# A behaviourally normed database of 1,377 natural sounds for auditory cognition and neuroscience

**DOI:** 10.64898/2026.08.25.746933

**Authors:** Marie Plegat, Maria Araújo Vitória, Giorgio Marinato, Beatrice Tita, Chris van der Lans, Minne Pijfers, Mia-Sara Bertovic, Michele Esposito, Elia Formisano, Bruno L. Giordano

**Affiliations:** Institut des Neurosciences de La Timone, UMR 7289, CNRS and Aix-Marseille University, Marseille, France; Department of Cognitive Neuroscience, Faculty of Psychology and Neuroscience, Maastricht University, Maastricht, Netherlands; BISS institute, Faculty of Science and Engineering, Maastricht University, Maastricht, The Netherlands

**Keywords:** Natural sounds, Auditory cognition, Behavioural norming, Sound event recognition, Computational modelling

## Abstract

Natural-sound research requires stimulus sets that combine acoustic standardization with detailed behavioural characterization. We present 1,377 two-second sounds representing 240 expert-defined source– action classes. We call this database “MaMa Sounds”, as it resulted from the collaborative effort of two academic teams in Maastricht and Marseille. The sounds were manually curated, segmented, sampled at 16 kHz, and labelled with a noun identifying the source and a verb identifying the action. We release deidentified trial-level identification and familiarity data together with multiple per-sound norms (e.g., identification accuracy, confidence and agreement; familiarity), along with overall norms derived with principal component analysis. Noun, verb, and joint noun–verb norms are provided as direct means and medians with the number of contributing observations. This battery preserves process-specific information, while two principal-component scores provide compact overall behavioural-identifiability measures derived from response ease, semantic correspondence, agreement, and familiarity. The repository also contains deterministic response-cleaning code, participant and reference Word2Vec representations, and code reproducing the public sound-level tables. The resource supports stimulus selection, matching, and continuous modelling in auditory cognition and neuroscience.

## 1 Background & Summary

Everyday listening requires the auditory system to infer sound-generating sources, actions, and events from complex acoustic signals. Natural sounds are therefore widely used to investigate how acoustical information is transformed into meaningful object and event representations in healthy listeners [15, 13, 4, 24, 20, 11], as well as in clinical and hearing-impaired populations [6, 27]. Their ecological richness is also a methodological challenge: experimental sounds must be controlled not only for duration and level, but also for recognizability, familiarity, ambiguity, and the relation between a source and the action or mechanism that produces the sound [19, 3, 2, 26].

Existing resources often trade breadth against experimental control. Large machine-listening datasets provide broad coverage of everyday events, but their clips may vary in duration, recording quality, background content, and number of concurrent sources, and they generally provide few behavioural norms from human listeners [8, 31]. Classic resources developed for behavioural or clinical research provide richer human norms but substantially smaller stimulus inventories. Marcell and colleagues reported naming, confidence, familiarity, complexity, pleasantness, response-latency, and categorization norms for 120 sounds; NESSTI provided multidimensional cognitive and affective norms for 110 standardized sounds; and Burns and Rajan related identification and subjective ratings to acoustic descriptors for 158 sounds [21, 16, 5]. More recently, the EnviSounds preprint introduced 530 sounds organized as 53 categories with ten exemplars each, together with recognition, response-latency, imageability, similarity, and goodness-of-exemplar measures [18]. Across this literature, identification has also been operationalized through different task formats, including free naming, sound-picture matching, and closed-set recognition [21, 25, 27]. A resource intended for behavioural and neuroimaging studies benefits from combining the scale of contemporary sound-event collections with manual curation, standardized waveforms, multiple complementary norms, and trial-level human data.

Semantic characterization poses an additional problem because natural sounds can be described at several levels. Broad categories such as “animal”, “human”, or “tool” are useful for some designs, but they omit the event-specific information listeners commonly report. Environmental-sound descriptions often identify both the sound-generating source and the action or mechanism, as in “dog barking” or “door closing” [28, 13, 12]. Source-action labels preserve this ecological specificity and allow source and action information to be represented separately or jointly.

Identifiability itself is also multidimensional. Previous norming studies have operationalized it through modal naming agreement, proportion correct, response latency, confidence, familiarity, and related perceptual ratings [2, 21, 13, 16, 5]. Ballas showed that identification time and accuracy covary with causal uncertainty and ecological experience, while later work used multivariate analyses to characterize shared structure among subjective and acoustic sound descriptors [2, 5]. These measures are related but not interchangeable. A sound may be recognized quickly yet receive several plausible labels; it may elicit strong cross-listener convergence without exactly matching an expert label; or it may be highly familiar but difficult to describe with one noun and one verb. Providing a battery of norms therefore permits process-specific stimulus selection and modelling. At the same time, shared variation among response speed, replay behaviour, confidence, semantic correspondence, agreement, and familiarity can be summarized in a compact overall score. Releasing both the component norms and principal-component summaries avoids forcing all users to adopt a single definition of identifiability. The dimensions assembled here have generally been distributed across smaller stimulus sets or different task formats; MaMa Sounds brings them together within one standardized inventory while retaining every component norm for separate use.

We created a database of 1,377 isolated natural sounds spanning 240 expert-defined noun-verb classes. We call this database “MaMa Sounds”, as it resulted from the collaborative effort of two academic teams in Maastricht and Marseille. The audio files included in MaMa Sounds are derived from clips distributed as part of FSD50K [8] and originally uploaded to Freesound [9]. The release combines standardized audio, detailed source-action labels, and labels for wide supra-ordinate categories, deidentified trial-level identification and familiarity data, participant and reference Word2Vec representations, and per-sound mean and median norms of identification response time, number of playbacks, confidence, reference similarity, reference-retrieval percentile, between-listener agreement, and familiarity. Two principal-component scores provide overall behavioural-identifiability measures, while the individual norms and participant-level data remain available for alternative participant-selection, text-processing, and aggregation choices. Individual audio files retain the Creative Commons license associated with their corresponding source clip. The resource is intended for stimulus selection, matching, and continuous modelling in behavioural, neuropsychological, computational, and neuroimaging studies of natural-sound recognition.

## 2 Methods

### 2.1 Sound selection and preprocessing

The selection and curation of sounds were carried out by six expert listeners. The collection was assembled from FSD50K [8], a dataset of 51,197 human-annotated Freesound [9] clips spanning 200 classes of the AudioSet ontology [10]. We initially selected up to six exemplars per source-action class, retaining recordings that clearly represented the intended event and excluding ambiguous, multi-event, or substantially contaminated recordings. Expert labels were standardized as a noun identifying the sound-generating source and a verb identifying the action or mechanism [13, 12]. When an acting agent and the sound-generating object differed, the object was preferred, as in “door closing” rather than “person closing”. For living sources, the noun could identify the organism when the sound was generated by its body, as in “dog barking”. The final collection comprised 1,377 sounds from 240 unique noun-verb classes.

Each recording was manually segmented from an identified event onset, trimmed or zero-padded to 2 s, demeaned, and level-normalized using the maximum root-mean-square amplitude measured in a 50-ms moving window. For 1,375 sounds, the peak 50-ms RMS differed by less than 0.0003 dB from the maximum database value. The two remaining scissor recordings differed by 0.185 and 0.386 dB. The released files are mono 16-bit PCM recordings sampled at 16 kHz and contain 32,000 samples each.

### 2.2 Semantic labels and clusters

Separate 300-dimensional Google News Word2Vec representations [22] were generated for the expert noun and verb labels using the same procedure applied to participant responses, described below. Unit-normalized noun and verb vectors were combined with equal weighting:

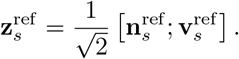

The 1,360 non-speech sounds were partitioned into 15 semantic clusters by applying k-means clustering with cosine distance to the joint noun–verb representations. The 17 speech sounds were excluded from this partition and assigned to a separate speech cluster, yielding 16 clusters in total. The choice of 15 non-speech clusters was pragmatic rather than an estimate of a privileged or “true” number of natural-sound categories. The clusters were used as relatively fine balancing strata when constructing behavioural sound sets; a finer partition was preferred to a coarser one to reduce the risk that narrower semantic concentrations remained unevenly distributed across sets. A short descriptive name was finally created for each cluster, based on the noun-verb description of the sounds they contained: `01 RingType`, Ringing and Typing Sounds (N sounds = 86); `02 NatuElem`, Nature and Weather Sounds (46); `03 HousWork`, Household and Work Sounds (127); `04 VehiEngi`, Vehicles and Engines (90); `05 DoorDraw`, Doors and Drawers (47); `06 BuzzWhir`, Buzzing and Whirring (120); `07 MechShoo`, Mechanical and Shooting (52); `08 MusiInst`, Musical Instruments (308); `09 LiquWate`, Liquid and Water Sounds (56); `10 TinkRust`, Tinkling and Rustling (44); `11 WarnAlar`, Warning/Alarm Sounds (58); `12 HumaVoca`, Human Vocal Sounds (115); `13 AnimBird`, Animal and Bird Sounds (84); `14 HumaNonv`, Non-vocal Human Sounds (85); `15 AnimDome`, Domestic Animal Sounds (42); and `16 Speech`, Speech (17). Note that these cluster labels are descriptive aids rather than claims that the partition constitutes a definitive taxonomy of natural sounds.

### 2.3 Behavioural experiments

Behavioural norms were collected in two online experiments with participants recruited through Prolific. The identification experiment was hosted on a server at Maastricht University, and the familiarity experiment was hosted on Pavlovia. Participants in both experiments were adult native English speakers. They were paid at an average hourly rate of 12.39 GBP, and completed the identification and familiarity experiments in 79.48 and 44.84 minutes, respectively (identification: SD = 28.73; median = 78.59; interquartile range IQR = 41.93 minutes; familiarity: SD = 15.20; median = 40.80; IQR = 15.01 minutes).

#### 2.3.1 Constrained verbal identification and confidence

The 1,377 sounds were partitioned into nine heterogeneous identification sets containing 150-156 sounds. Semantic context can affect environmental-sound identification [3, 27, 14]; the sets were therefore designed to remain semantically heterogeneous. Four sets had been defined for companion neuroimaging studies [23, 30, 1], and the remaining sounds were assigned while balancing the 16 semantic clusters as closely as possible. Set membership is included in the released metadata.

On every trial, participants could replay the sound and responded to the prompts “Please describe the sound source (noun)” and “Please describe the sound action (verb)”. They then rated confidence in the identification on a 0-10 scale. The number of sound presentations was retained as the identification number of playbacks.

Identification response time was derived from the positive difference between consecutive values of the cumulative active-task timer. The first trial of each session was coded as missing because no explicit timestamp for the onset of the first sound was available. Non-positive differences and timer resets were also coded as missing. This measure includes listening, replaying, identification, typing, and confidence reporting and is therefore a trial-level identification response time rather than a pure perceptual reaction time.

The raw exports contained 426 candidate runs. Before participant-level quality-check (QC), 104 confidence values outside the valid 0-10 range were coded as missing; no confidence values were rescaled. A run passed structural QC when no more than 15 expected trials were incomplete and no major filename or identification-set corruption was detected. An expected trial was incomplete when the noun response, verb response, or confidence value was missing. Two otherwise structurally compliant runs were excluded because more than 50% of the non-missing responses in at least one verbal field were grammatical placeholders. Sixty runs failed one or more QC criteria. When a Prolific ID had more than one QC-compliant run, the earliest QC-compliant run was retained and later runs were excluded, including runs involving a different identification set. One later QC-compliant duplicate run was excluded.

The final identification sample comprised 365 unique participants, with 37-44 participants per identification set. As a whole, these participants contributed a total of 55,480 complete identification trials. Age was available for 356 participants (mean = 27.52 years, SD = 4.56, median = 28, IQR = 8, range = 18–35). The sample included 168 participants recorded as female, 185 as male, and 12 classified as other under the reporting scheme.

#### 2.3.2 Familiarity ratings

The 1,377 sounds were partitioned into seven approximately cluster-balanced rating sets containing 196 or 197 sounds. Participants rated each sound twice on a 0-100 familiarity scale, once in each half of the experiment. The scale endpoints were labelled “Not familiar” and “Very familiar”. Before the rating task, participants completed a modified version for the JsPsych framework [7], of the antiphase-based headphone-screening procedure of Woods et al. [32], with up to two attempts.

The raw exports contained 298 candidate submissions. A submission passed QC only when the headphone check was explicitly passed, the Pearson correlation between the two rating repetitions was at least 0.10, and the participant used at least 10 distinct valid rating values. Four submissions failed the headphone check, four had a missing or invalid headphone-check result, 12 failed the test-retest criterion, and 10 failed the rating-diversity criterion; these criteria overlapped, yielding 19 QC-noncompliant submissions in total. Of the 279 QC-compliant submissions, four later submissions from repeated Prolific IDs were excluded, retaining the earliest QC-compliant submission.

The final familiarity sample comprised 275 unique participants, with 32-52 participants per rating set. The included submissions contributed 108,212 valid familiarity-rating trials across the two repetitions. Age was available for 260 participants (mean = 36.12 years, SD = 9.95, median = 35, IQR = 13, range = 18-59). Gender information was recorded as female for 116 participants, male for 144, another response category for 10, and missing for 5.

### 2.4 Ethics

The experiments were approved by the Ethics Review Committee Pusychology and Neuroscience (ERCPN) at Maastricht University (approval OZL 239 106 06 2021 S2) and the Aix-Marseille University ethics committee (approval 0062025-39). All participants provided informed consent before taking part, including consent for sharing deidentified research data.

### 2.5 Processing of verbal identifications

The original noun and verb responses were retained and were not manually recoded. A deterministic pipeline applied Unicode normalization, case folding, white space normalization, boundary-punctuation normalization, and removal of predefined prompt or discourse wrappers. Technical missing values, explicit statements of uncertainty, grammatical placeholders, responses without alphabetic content, and responses empty after wrapper removal were marked invalid. The response recoding procedure did not apply spelling correction, lemmatization, translation, semantic rewriting, noun-verb field swapping, or imputation.

Valid cleaned responses were represented using the 300-dimensional Google News Word2Vec model [22]. A deliberately small set of non-informative function words was removed, including articles, infinitival markers, conjunctions, and auxiliaries such as `a`, `an`, `the`, `to`, `of`, `and`, `or`, `is`, `are`, `was`, `were`, `be`, `been`, and `being`; the complete list is provided in the released code. Exact vocabulary keys and supported underscore compounds were then identified. Each selected key vector was normalized to unit length so that every retained key contributed equal weight, the available vectors were averaged, and the resulting response vector was normalized to unit length. The same procedure was applied to the expert noun and verb labels. Responses without a supported vocabulary key were treated as unavailable rather than imputed.

Noun and verb responses were represented independently (noun or verb embedding) or, in separate analyses, jointly. For trials with both representations, the joint vector was defined as

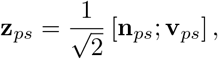

where **n**_*ps*_ and **v**_*ps*_ are unit-normalized noun and verb vectors for participant *p* and sound *s*. This construction gives equal weight to source and action information and yields a unit-normalized joint vector.

### 2.6 Sound-level behavioural norms

#### 2.6.1 Reference similarity and reference retrieval

Reference similarity was the cosine similarity between a participant response vector and the corresponding expert-label vector. It was computed separately for nouns, verbs, and joint noun-verb vectors. Larger values indicate greater Word2Vec proximity to the expert label.

Reference-retrieval percentile provided an inventory-relative measure of semantic correspondence. For each participant response, its similarity to the correct expert reference was compared with its similarities to every unique nonmatching expert reference in the same response domain. Noun candidates were the 175 unique noun labels, verb candidates were the 138 unique verb labels, and joint candidates were the 240 unique noun-verb classes observed in the database. Exact duplicate labels were represented once, and the correct target was removed from the comparison set.

Let 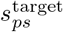 denote the cosine similarity between the response of participant *p* to sound *s* and its correct reference, and let 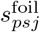 denote its similarity to nonmatching candidate *j*. The retrieval percentile was

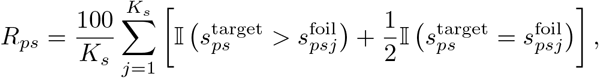

where *Ks* is the number of nonmatching candidates and ties receive half credit. A value of 50 indicates that the target outranked half of the nonmatching references, whereas 100 indicates that it outranked all alternatives. With half credit for ties, the statistic has a probability-of-superiority interpretation analogous to rank-based common-language measures [29], although its candidate inventory and retrieval interpretation are specific to MaMa Sounds. It is therefore a reference-inventory retrieval percentile rather than a human-calibrated percentage correct.

#### 2.6.2 Between-listener agreement

Between-listener agreement was defined from the cosine similarities between all unordered pairs of participant response vectors for a sound. For *Ns* available response vectors, mean agreement was

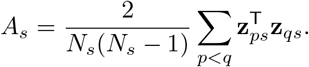

with *p* and *q* indexing the response vectors of different participants. The direct median of the unordered pairwise similarities was also retained. Agreement was calculated separately for noun, verb, and joint noun-verb representations and was undefined when fewer than two response vectors were available. The associated `n used` field reports the number of participant vectors, not the number of pairs.

#### 2.6.3 Aggregation and missing values

For every sound, direct means and medians were calculated from all available eligible observations for identification response time, number of playbacks, confidence, reference similarity, reference-retrieval percentile, and agreement. Missing values were handled independently for each norm family and were not imputed. A valid noun response could therefore contribute to noun norms when the verb was unavailable, whereas joint norms required both representations. The released table reports the number of contributing observations for every norm family.

For familiarity, all available valid repetitions were first averaged within participant and sound. These participant-specific values were then averaged or median-aggregated across participants. A single valid repetition was retained when its paired repetition was missing. Familiarity ratings were not imputed.

### 2.7 Overall behavioural-identifiability norms

Two principal-component scores summarized shared variation across six sound-level norms: identification response time, identification number of playbacks, identification confidence, joint noun-verb reference-retrieval percentile, joint noun-verb agreement, and familiarity. For the mean-derived score, the six mean norms were z-scored using the population standard deviation. For the median-derived score, average tied ranks were calculated for each median norm and then z-scored. Identification response time and number of playbacks were sign-reversed so that larger values consistently indicated easier identification.

Principal-component analysis [17] was fitted separately to the two six-variable matrices using full singular-value decomposition without whitening. The first component was sign-oriented to correlate positively with the mean of the six direction-aligned inputs. The released PC1 scores were then re-standardized to mean zero and unit population standard deviation. The individual component norms remain available and should be used when a process-specific measure is required.

## 3 Data Records

The complete public release is deposited at Zenodo (doi:10.5281/zenodo.22042882). The release is distributed as a ZIP archive; the directory structure below refers to its contents after extraction. All released tabular files are UTF-8 tab-separated-value files.

- `data/sounds/` contains the 1,377 mono 16-bit PCM WAV files. Each sound is 2 s long, sampled at 16 kHz, and contains 32,000 samples.
- `data/identification/` contains 365 deidentified participant-level identification files. Each row contains the trial index, sound filename, number of playbacks, original noun response, original verb response, confidence value, and identification response time.
- `data/identification/01_automated_cleanup/` contains the original identification rows with appended deterministic cleaned noun and verb fields and Boolean validity flags.
- `data/identification/02_w2v/` contains the cleaned participant responses with the exact Google News keys used, separate 300-dimensional noun and verb vectors, availability flags, and vocabulary-coverage values.
- `data/familiarity_ratings/` contains 275 deidentified participant-level familiarity files. Each row contains a consecutive trial number, sound filename, rating repetition, and familiarity rating.
- `data/MaMA_Sounds_reference_w2v_embeddings.tsv` contains the expert noun- and verb-label Word2Vec representations generated using the same procedure as the participant responses.
- `data/MaMa_Sounds_table.tsv` is the canonical 1,377-row master table. It contains 13 stimulus-metadata fields and 41 behavioural norm fields. The metadata fields are `MaMa idx`, `wav name`, `ident set`, `rating set`, `MaMa cluster`, `MaMa label noun`, `MaMa label verb`, `FSD50K class`, `FSD50K filename`, `general category`, `fs`, `n samples`, and `peak rms 50ms dbfs`. The 41 norm fields comprise direct mean norms, direct median norms, the corresponding `n used` values, and two overall PC1 norms.
- `data/MaMa_Sounds_overall_norm_pca_parameters.tsv` records the PCA input variables, preprocessing centers and scales, direction multipliers, PC1 weights, input correlations, explained variance, and software version.
- `data/MaMa_Sounds_table_correlations_mean_pearson.tsv` and `data/MaMa_Sounds_table_correlati ons_median_spearman.tsv` contain the 14-by-14 across-sound validation matrices. The corresponding annotated PNG files are included under the same filenames with `correlation matrix` in place of `correlations`.
- `code/` contains four public processing scripts and one shared response-cleaning helper. These scripts regenerate deterministic cleanup, participant and reference Word2Vec representations, sound-level norms, PCA scores, and validation outputs from the deposited deidentified participant files. The Google News Word2Vec model weights are not redistributed.

Public participant identifiers are experiment-specific and are not cross-linked between identification and familiarity. Original platform exports, direct identifiers, demographics, private QC tables, excluded submissions, and private tests are not deposited. Each `n used` field reports the number of participant observations or vectors contributing to the corresponding mean and median norms.

## 4 Technical Validation

### 4.1 File integrity and observation coverage

The released audio and metadata were checked for one-to-one filename correspondence. All 1,377 files were mono 16-bit PCM recordings sampled at 16 kHz and contained 32,000 samples. The final master table contained one row for every sound. No behavioural value or unavailable embedding was imputed.

The included identification sample contributed 55,480 complete trials, and the included familiarity sample contributed 108,212 valid rating trials. Across sounds, 34-44 observations contributed to identification response time, 37-44 to number of playbacks, 33-44 to confidence and noun-based semantic norms, 32-44 to verb-based semantic norms, 31-44 to joint noun-verb semantic norms, and 32-52 participants to familiarity.

### 4.2 Reference retrieval

The sounds were generally well identified relative to the MaMa Sounds reference inventory. Across sounds, the median participant-mean reference-retrieval percentile was 88.0 for nouns (IQR = 21.1), 82.9 for verbs (IQR = 21.9), and 89.9 for joint noun-verb representations (IQR = 16.7). The corresponding participant-median norms had across-sound medians of 96.6, 94.5, and 97.1, respectively (Fig. 1).

**Figure 1.**
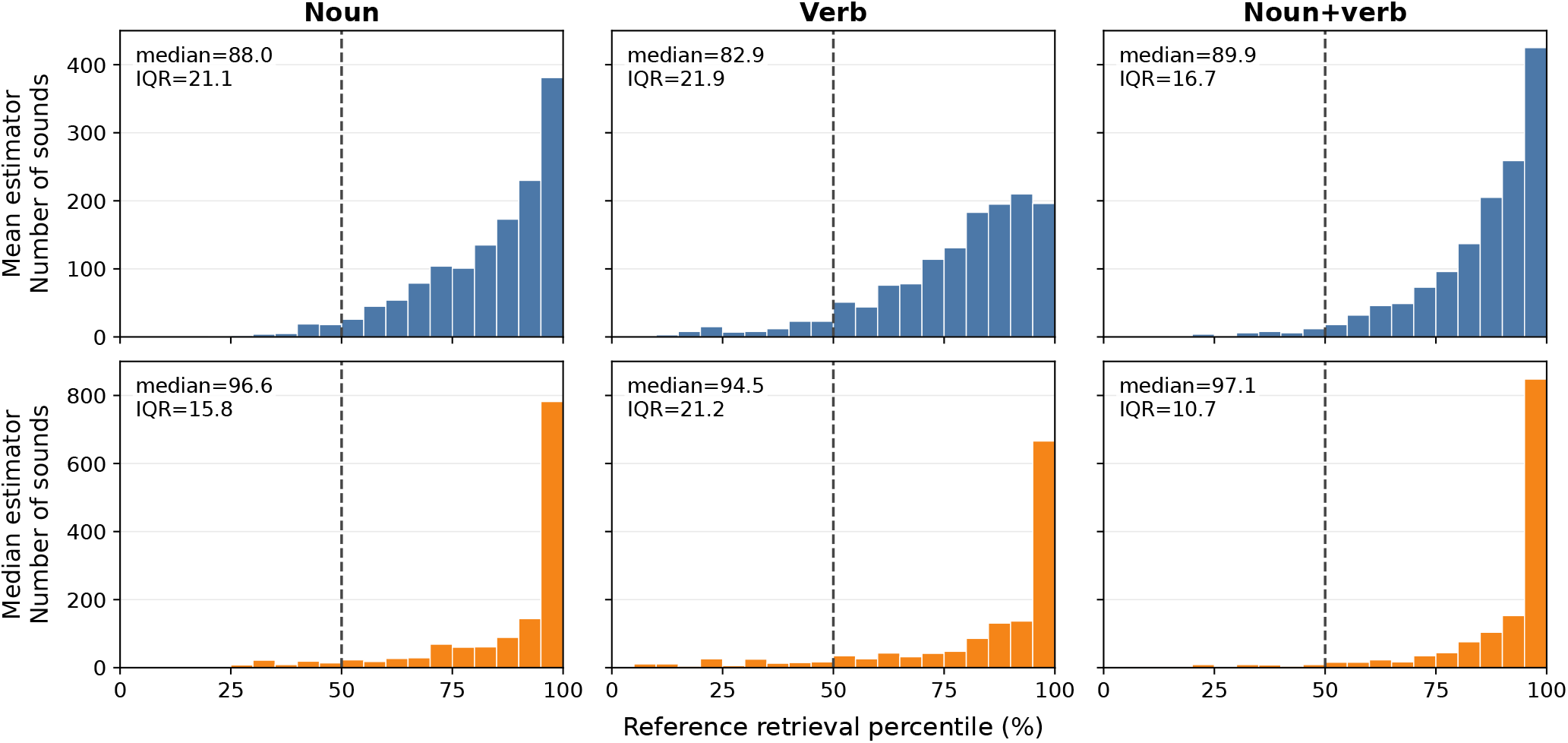
Reference-retrieval percentiles across the 1,377 sounds. Histograms show the distributions of sound-level noun, verb, and joint noun-verb retrieval percentiles. The upper row shows means across eligible participant responses; the lower row shows medians. The dashed vertical line at 50 marks the expected percentile when the correct reference outranks half of the nonmatching candidate inventory. Panel annotations report the across-sound median and interquartile range.

### 4.3 Consistency among behavioural norms

The mean joint retrieval percentile correlated positively with identification confidence (*r* = .631), joint agreement (*r* = .669), familiarity (*r* = .495), and raw joint reference similarity (*r* = .754), and negatively with identification response time (*r* = −.278) and number of playbacks (*r* = −.563). The median-derived Spearman correlations showed the same pattern (*ρ* = .641, .593, .562, .755, −.315, and −.462, respectively). The full mean-derived correlation matrix is shown in Fig. 2; correlations involving the overall norm describe the composition of the PCA score and are not independent validation of that score.

**Figure 2.**
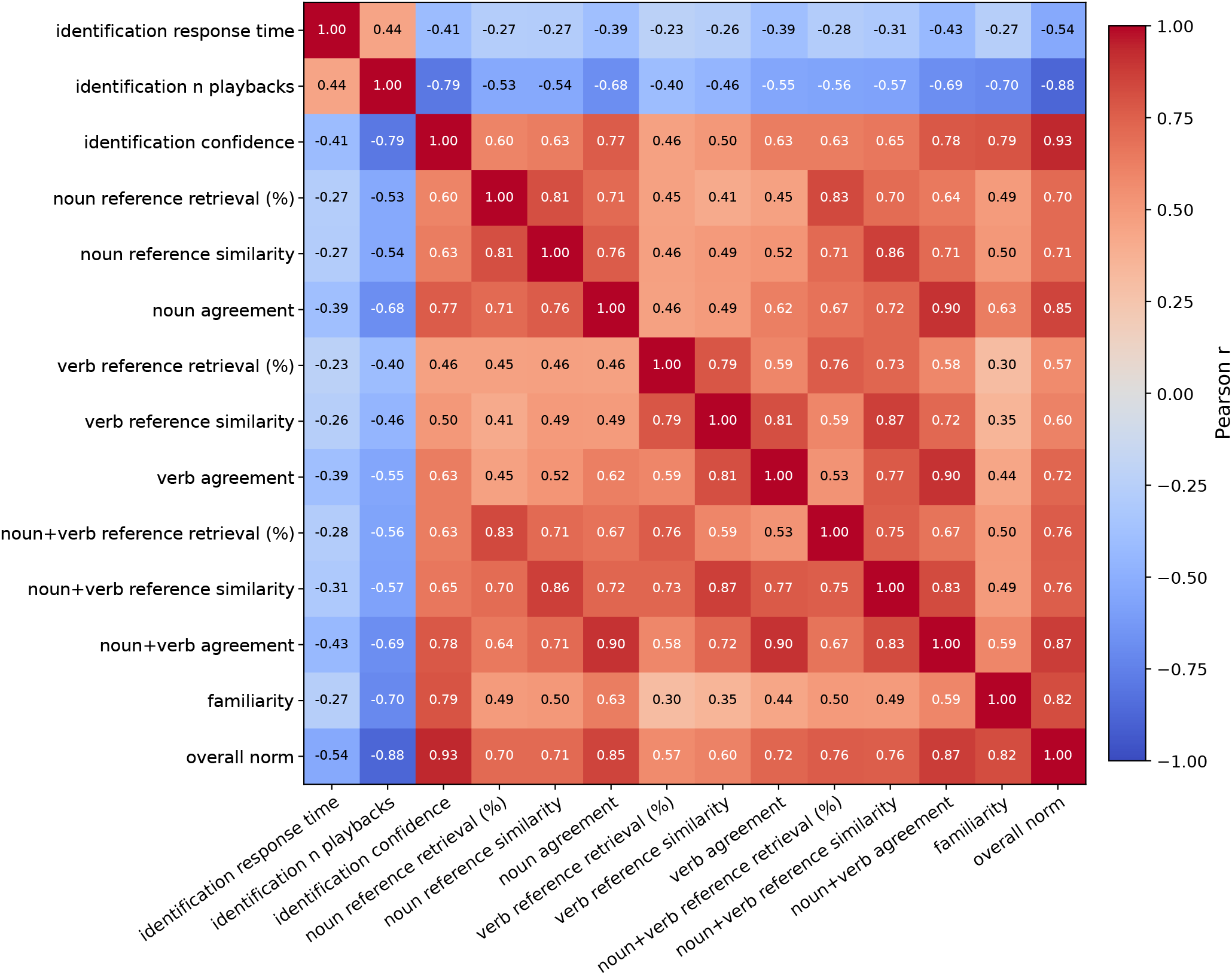
Across-sound associations among mean-derived behavioural norms. Cells show Pearson cor-relations across the 1,377 sounds for identification response time, number of playbacks, confidence, noun, verb, and joint reference-retrieval percentile, noun, verb, and joint reference similarity, noun, verb, and joint agreement, familiarity, and the mean-derived overall PC1 norm. The overall norm is computed from response time, number of playbacks, confidence, joint retrieval percentile, joint agreement, and familiarity; its correlations with those inputs therefore characterize the composite rather than provide independent validation.

### 4.4 Familiarity consistency

Among the 275 included participants, the mean participant-level correlation between the two familiarity repetitions was .647 and the median was .706 (range = .107–.925). The complete retained sample contained 108,212 valid ratings across the two repetitions. These values support the use of participant-averaged familiarity ratings as sound-level norms.

### 4.5 Overall behavioural-identifiability norms

The first principal component explained 65.4% of the variance in the mean-derived six-norm matrix and 63.7% in the median-rank-derived matrix (correlation between the two PC scores = .967. These scores provide compact summaries of shared behavioural variation, while all component norms remain available for process-specific analyses.

## 5 Usage Notes

Identification response time is not a perceptual reaction time. It includes sound presentation, possible replay, identification, typing, confidence reporting, and other within-trial activity. The first trial of every identification session has no response-time estimate.

All waveforms are sampled at 16 kHz, so their Nyquist frequency is 8 kHz. The resource is therefore not intended for analyses requiring natural acoustic content above 8 kHz. Waveform-level RMS normalization standardizes the digital stimuli, but it does not imply that playback sound-pressure level was calibrated across participants’ online listening environments.

Word2Vec reference similarity measures absolute proximity to the expert label in the specified embedding space. Reference-retrieval percentile measures how strongly the intended label is retrieved relative to the other MaMa Sounds expert labels. It is defined relative to the labels part of the released dataset, and should not be interpreted as a human-calibrated percentage correct. Agreement measures semantic convergence across listeners independently of the expert label.

Mean norms retain the contribution of all eligible participant responses and are more sensitive to minority alternative identifications. Median norms describe the central participant response and are more robust to extreme values, but several median retrieval norms approach the upper end of the scale. Users should select the estimator appropriate to their application and consult the corresponding `n used` field.

The overall PC1 norms summarize shared variation in response speed, replay behaviour, confidence, semantic retrieval, agreement, and familiarity. They are broad behavioural-identifiability measures rather than pure accuracy scales. Users interested in a specific process should use the corresponding component norm.

Word2Vec-derived norms are conditional on the Google News vocabulary and the released preprocessing choices. Responses without any supported key are treated as unavailable rather than imputed. Because free naming and the Google News model are English-language dependent, naming-derived semantic norms may not transfer unchanged across linguistic or cultural populations. The deposited participant-level text and vectors permit alternative participant-inclusion rules, text-processing procedures, semantic models, and aggregation strategies.

## 6 Data Availability

The complete public release, including the 1,377 sound files, deidentified participant-level identification and familiarity data, participant and reference Word2Vec representations, final per-sound norms, PCA parameters, and validation tables, is available from Zenodo at doi:10.5281/zenodo.22042882. The repository record and README describe the ZIP archive contents, folder structure, and file formats.

## 7 Code Availability

The custom Python code used to generate deterministic response cleanup, participant and reference Word2Vec representations, per-sound behavioural norms, principal-component scores, and validation tables from the deposited deidentified participant files is included in the same repository under `code/`. The Google News Word2Vec model weights are not redistributed; the required model filename and SHA-256 checksum are documented in the repository README. Reconstruction from the original platform exports, deidentification, and participant-level QC used restricted identifiers and are not part of the public processing workflow.

## Author Contributions

Conceptualization: BLG, EF. Experiment design: BLG, GM, CvdL. Data acquisition: GM, MAV, BT, CvdL, MSB. Data curation: BLG, GM, EF, MPl, MAV,ME, MPi. Formal analysis: BLG, GM. Funding acquisition: BLG, EF. Investigation: MPl, MAV, GM, BT, CvdL, MPi, MSB, EF, BLG. Methodology: BLG, EF, MPl, MAV, GM, ME, CvdL. Project administration: BLG, ME, EF. Resources: BLG, GM, MPl, EF, CvdL. Software: BLG, EF, GM, CvdL. Supervision: BLG, EF. Validation: BLG, MPl, GM, ME. Visualization: BLG, GM. Writing – original draft: MPl, BLG, GM. Writing – review and editing: BLG, MPl, GM, EF, ME, MAV.

